# T cells instruct dendritic cells to produce inflammasome independent IL-1β causing systemic inflammation

**DOI:** 10.1101/475517

**Authors:** Aakanksha Jain, Ricardo A. Irizarry-Caro, Amanpreet S. Chawla, Naomi H. Philip, Kaitlin R. Carroll, Jonathan D. Katz, Andrew Oberst, Alexander V. Chervonsky, Chandrashekhar Pasare

**Affiliations:** Immunology Graduate Program, University of Texas Southwestern Medical Center at Dallas, TX, 75390, USA; Division of Immunobiology, Children’s Hospital Medical Center, Cincinnati, OH, 45225, USA; Center for Inflammation and Tolerance, Cincinnati Children’s Hospital Medical Center, Cincinnati, OH, 45225, USA; Department of Pathobiology, School of Veterinary Medicine, University of Pennsylvania, Philadelphia, PA, 19104, USA; Department of Immunology, University of Washington, Seattle, WA, 98109, USA; Department of Pathology, University of Chicago, Chicago, IL, 60637, USA

**Author notes:** Correspondence to Chandrashekhar Pasare.

## Abstract

While IL-1β is critical for anti-microbial host defense, it is also a key mediator of autoimmune inflammation. Inflammasome activation following pathogenic insults is known to result in IL-1β production. However, the molecular events that produce IL-1β during T cell driven autoimmune diseases remain unclear. Here, we have discovered an inflammasome-independent pathway of IL-1β production that is triggered upon cognate interactions between dendritic cells and effector CD4 T cells. Analogous to inflammasome activation, this “T cell-instructed IL-1β also relies on two independent signaling events. TNFα produced by activated CD4 T cells engages TNFR signaling on DCs leading to pro-IL-1β synthesis. Subsequently, FasL, also expressed by effector CD4 T cells, engages Fas on DCs leading to caspase-8 dependent pro-IL-1β cleavage. Remarkably, this two-step mechanism is completely independent of pattern recognition receptor activation. IL-1β produced upon cognate DC-effector CD4 T cell interaction causes wide spread leukocyte infiltration, a hallmark of systemic inflammation as well as autoimmune pathology. This study has uncovered a novel feature of DC-T cell cross-talk that allows for active IL-1β secretion independent of innate sensing pathways and provides a mechanistic explanation for IL-1β production and its downstream consequences in CD4 T cell driven autoimmune pathology.

---

IL-1β mediates host immunity through its ability to influence both innate and adaptive immune responses. It promotes innate immunity by inducing the acute phase response and recruiting inflammatory cells^1–4^. In the adaptive immune system, IL-1β enhances T cell priming and differentiation^5^, and more importantly acts as a licensing cytokine to enable memory CD4 T cell effector function^6^. However, aberrant production of IL-1β in the absence of pathogenic insult can result in immunopathology associated with several auto-immune and auto-inflammatory diseases^7^. In fact, IL-1β blockade has been proven to be a successful treatment for several autoimmune conditions including psoriasis, rheumatoid arthritis (RA), and uveitis^8^. Autoinflammatory disorders can be a consequence of gain-of-function mutations in inflammasome machinery which results in spontaneous IL-1β production^9–13^. Due to this association, IL-1β production in these disorders is often attributed to inflammasome activation^14^. In the case of T cell driven auto-immunity, where IL-1β is implicated as a major driver of pathology, the role of inflammasome in IL-1β production remains obsure^14–17^. GWAS studies have also failed to report significant genetic association between inflammasome proteins and T cell dependent autoimmunity^14^. Additionally, disease progression in mouse models of RA was found to be independent of the inflammasome components NLRP3 and caspase-1^16^. Similarly, caspase-1 deficiency did not mitigate diabetes in NOD mice^18^. Together, this prompted us to investigate how bioactive IL-1β is produced during T cell-driven autoimmune diseases in the absence of overt infection or injury.

We have previously demonstrated that T cell intrinsic IL-1R signaling is critical for optimal cytokine production by effector and memory CD4 T cells following their reactivation using DCs^6^. The DCs used in these experiments were not stimulated by any microbial ligands, thus leading us to hypothesize that cognate interaction between DCs and effector CD4 T cells might elicit the production of bioactive IL-1β independent of pattern recognition receptor (PRR) signaling. Indeed, cognate interaction of DCs with effector CD4 T cells led to secretion of IL-1β in the absence of innate immune sensing (Fig. 1a). Furthermore, we found that DC interaction with effector CD4 T cells of Th1, Th2 and Th17 lineages all led to secretion of IL-1β, pointing to a broadly conserved pathway of IL-1β production (Fig. 1b). Since all CD4 T cell lineages were found to instruct IL-1β production by DCs, we henceforth use Th0 cells to represent effector CD4 T cells, unless otherwise noted. These initial experiments were performed using anti-TCR antibody stimulation that activates T cells regardless of their specificity. We therefore examined if this IL-1β production was in fact dependent on presentation of cognate peptide by DCs. When OVA peptide-restricted OT-II TCR transgenic effector T cells were reactivated using DCs, IL-1β production was strictly dependent on the presence of the cognate OVA_323-339_ peptide (Fig. 1c). More interestingly, the quantity of IL-1β found in the supernatants directly correlated with the concentration of the stimulating peptide (Fig. 1d), indicating that the avidity of MHC-TCR interaction is a major determinant of the amount of IL-1β being secreted. While it has been reported that CD4 T cells can produce IL-1β^19^, we found that DCs were the only source of IL-1β during their interaction with effector CD4 T cells (Fig. 1e).

**Figure 1.**
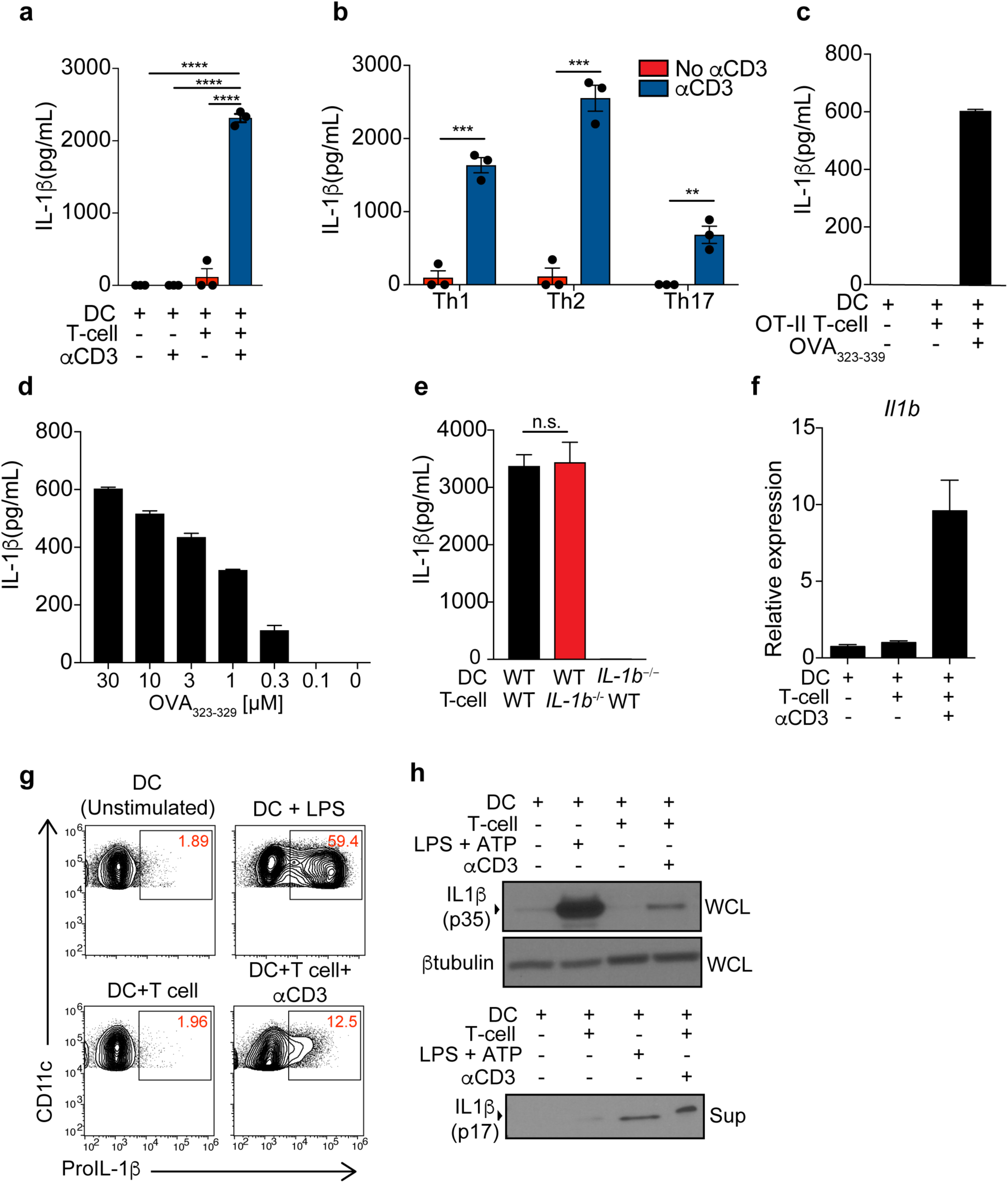
Cognate interaction between DCs and effector CD4 T cells leads to inflammasome independent production of bioactive IL-1β by DCs. (a) WT effector CD4 (Th0) cells were stimulated with WT DCs using αCD3 for a period of 18h and lL-1β was quantified in the supernatants. (b) Effector CD4 T cells polarized to Th1, Th2 and Th17 lineage were stimulated with WT DCs using αCD3 for 18h followed by IL-1β ELISA (c and d) IL-1β was quantified in the supernatants following stimulation of Th17 polarized OT-II T cells (18h) using (c) fixed (100µM) or (d) titrating concentrations of OT-II323-339 peptide presented by DCs (e) Effector CD4 (Th0) cells of given genotypes were stimulated with αCD3 using DCs of indicated genotypes and IL-1β was measured following 18h of culture (f) WT DCs and αCD3 were used to stimulate *Il1b^-/-^* Th17 cells for 3h and lysates from co-cultures were used to make cDNA to assess *Il1b* mRNA by qPCR. Data are normalized to 18s rRNA. (g) WT DCs were stimulated with LPS (100ng ml^-1^) or cultured with Th17 cells in the presence of αCD3 and brefeldin A for 6h. Intracellular pro-IL-1β expression in live CD90-ve CD11c+ve cells was measured using flow cytometry. (h) WT DCs were used to stimulate effector CD4 T cells (Th0) with αCD3 for 18h and pro-IL-1β (p35) and cleaved IL-1β (p17) levels were analyzed in the cell lysates and the supernatants, respectively, using Western blotting. Error bars indicate SEM; (a, b and e) paired *t*-test, (c,d,f-h) Data are representative of three independent experiments.

Production of IL-1β relies on two independent signals: a priming signal necessary for transcription and synthesis of proIL-1β, followed by a cleavage signal required for its biological activity^20^. We observed that DC-T cell interaction led to rapid transcriptional upregulation of *Il1b* (Fig. 1f). This experiment was performed using *Il1b^-/-^* T cells to ensure analysis of *Il1b* transcript specifically in DCs. The transcriptional induction translated into accumulation of intracellular pro-IL-1β (Fig. 1g). In addition to proIL-1β synthesis, interaction with effector CD4 T cells also induced cleavage of pro-IL-1β into its bioactive 17kDa fragment (Fig. 1h), which could be detected in the supernatant. These data demonstrate that interaction of DCs with effector CD4 T cells can trigger transcriptional induction of pro-IL-1β as well as its bioactive cleavage.

Next, we decided to characterize the signaling events that enable this IL-1β production. Inflammasome activation is a major mechanism by which IL-1β is produced^21^. The priming signal during inflammasome activation is provided by sensing of microbial ligands by pattern recognition receptors (PRRs) such as TLRs that culminates into NF-κB activation and transcriptional upregulation of pro-IL-1β^22^. Since DCs used in these experiments were not exposed to any microbial ligands, we posited that a PRR-independent signal is likely to be involved in proIL-1β synthesis. A significant difference in the magnitude of pro-IL-1β synthesis following TLR stimulation as compared to T cell interaction also suggested a TLR independent mechanism (Fig 1g). Nevertheless, to rule out the contribution of inadvertent endotoxin contamination in these *in vitro* cultures, we analyzed induction of proIL-1β in *Tlr2/4^-/-^* as well as *Myd88^-/-^* DCs. We found no defect in effector CD4 T cell driven IL-1β induction in DCs deficient for TLR signaling (Fig 2a and Supplementary Fig. 1a and 1b). It is well known that pro-IL-1β transcription is mediated by NFκB^22^ and AP-1^23^, which can also be activated downstream of receptors for the TNF superfamily. This prompted us to examine a potential role for these receptors in pro-IL-1β synthesis in the absence of microbial recognition by DCs. We directly tested the role of TNFR superfamily signaling in IL-1β production by neutralizing individual ligands during DC-T cell interaction. We found that neutralization of TNFα and FasL, but not CD40L, significantly compromised IL-1β production (Fig. 2b). This suggested that TNFR as well as Fas signaling participate in PRR independent IL-1β production by DCs. Upon careful examination, we observed that the pro-IL-1β levels were compromised only in the absence of TNFα (Fig. 2c), while Fas signaling was dispensable for pro-IL-1β synthesis (Fig. 2c). This result suggests that while Fas-FasL interaction is critical for bioactive IL-1β production, it is not required for the transcriptional induction of pro-IL-1β. Consistently, intracellular pro-IL-1β was also significantly reduced upon neutralization of TNFα (Fig. 2d). Moreover, stimulation of DCs with recombinant TNFα was sufficient to drive the synthesis of pro-IL-1β (Supplementary Fig. 1c). Since TNFα neutralization does not result in complete abrogation of pro-IL-1β, additional pathways could contribute to pro-IL-1β synthesis that require further investigation. The existence of diversified mechanisms of T cell induced pro-IL1β, is parallel to the ability of several innate recognition receptors to upregulate IL-1β^21,24,25^. We found that effector CD4 T cells of all lineages (Th1, Th2 and Th17) rapidly upregulate TNFα following their interaction with DCs (Supplementary Fig. 1d). While TNFα is primarily known to be an effector cytokine of the Th17 lineage^26^, we found that primed Th1 and Th2 lineage cells are also capable of producing TNFα (Supplementary Fig. 1d), suggesting that activated CD4 T cells of all lineages are poised to engage TNFR on DCs. It is important to note that while TNFα is known to be largely of myeloid origin in PAMP driven inflammation, activated T cells seem to be the predominant TNFα producers during their interactions with DCs (Fig. 2e and Supplementary Fig. 1e). Consistent with this finding, TNFα deficient T cells induced significantly diminished pro-IL-1β (Fig 2f) production by DCs. Finally, combined deficiency of TNFR1 and TNFR2 on DCs showed that DC intrinsic TNFR signaling was required for optimal IL-1β production (Fig. 2g).

**Figure 2.**
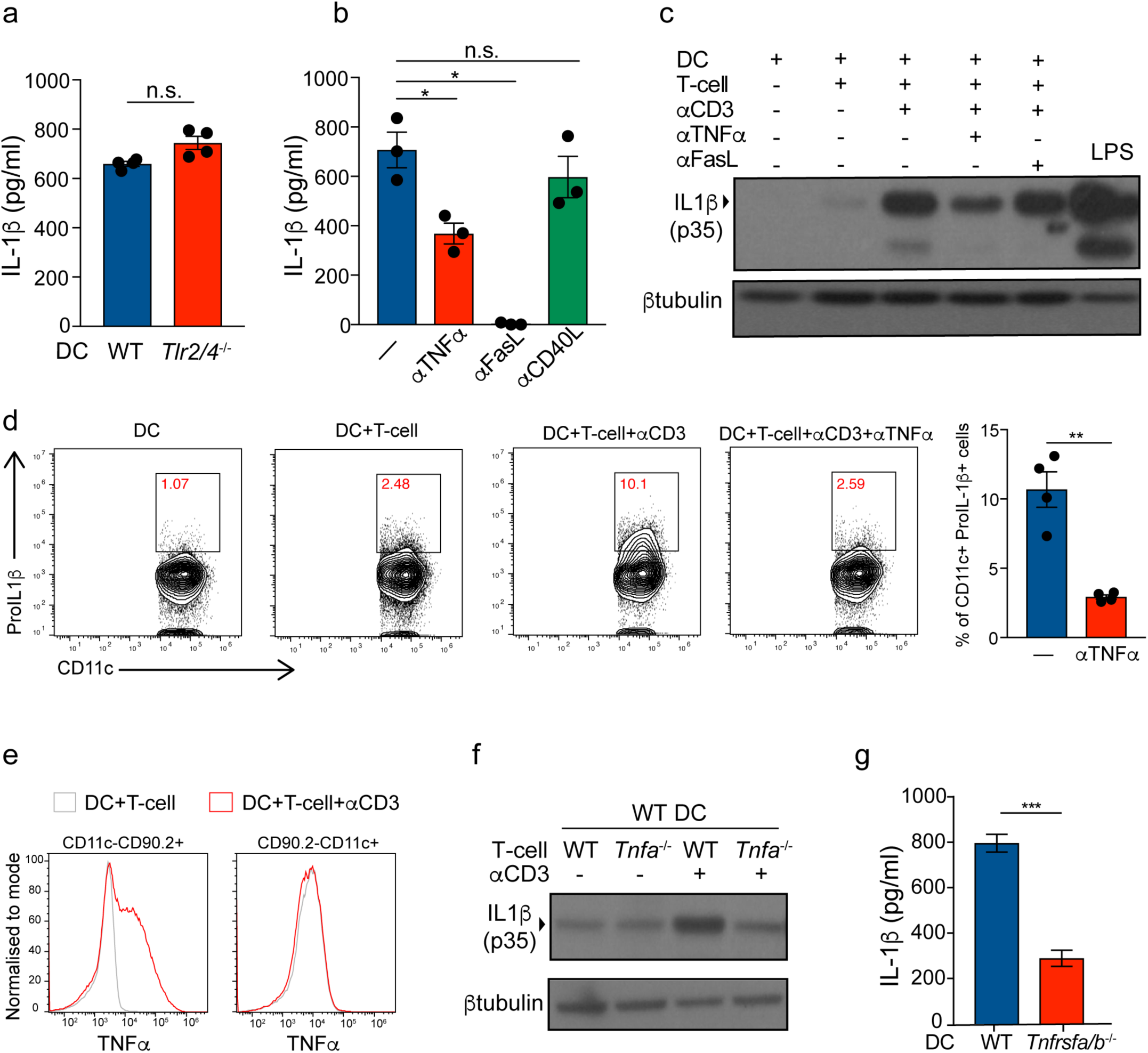
T cell derived TNFα is critical for induction of pro-IL-1β in DCs. (a) WT and *Tlr2/4^-/-^* DCs were stimulated with WT Th0 cells using αCD3. Secreted IL-1β was quantified 6h post stimulation (b) WT Th0 effector cells were stimulated with αCD3 and WT DCs in the presence or absence of anti-TNFα (20µg/ml), anti-FasL (10µg/ml), and anti-CD40L (20µg/ml), neutralizing antibodies. Secreted IL-1β in the culture supernatant was quantified after 6h of culture (c) Whole cell lysates from DCs cultured as indicated with effector CD4 T cells (Th0) in the presence or absence of antibodies were subject to Western blot analysis to detect pro-IL-1β. DCs stimulated with LPS were used as positive control to detect pro-IL-1β (d) Effector CD4 T cells (Th0) were stimulated with WT DCs using αCD3 in the presence or absence of anti-TNFα (20ug ml^-1^) neutralizing antibody for 6h. Intracellular pro-IL-1β was quantified using flow cytometry (e) Th0 cells were stimulated with αCD3 using WT DCs in the presence of brefeldin A for 3h. Intracellular TNFα in T cells and DCs was quantified using flow cytometry. (f) WT or *Tnfa^-/-^* DCs were used to stimulate WT or *Tnfa^-/-^* effector CD4 T cells and Western blot was performed on whole cell lysates (WCL) following 6h after αCD3 stimulation. (g) WT or *Tnfra/b*^-/-^ DCs were used to stimulate WT effector CD4 T cells (Th0) for 6h in the presence of αCD3 and supernatants were collected for IL-1β ELISA. Error bars indicate SEM; (a,b,d,g) paired *t*-test, (c,d,e,f) Data are representative of two independent experiments.

We then proceeded to identify the molecular mechanism responsible for proteolytic cleavage of pro-IL-1β. Caspase-1, the effector protease of all inflammasomes^27^, is largely responsible for bioactive cleavage of pro-IL-1β^14^. Indeed, we found that the aspartate residue in pro-IL-1β, where caspase-1 cleaves, is critical for IL-1β production following DC-T cell interaction (Supplementary Fig. 2a). Surprisingly, we did not detect active caspase-1 following DC-T cell interaction (Fig. 3a). Furthermore, we found that genetic ablation of caspase-1 in DCs did not compromise IL-1β release (Fig. 3b and Supplementary Fig. 2b). These data point to the existence of an inflammasome independent mechanism of IL-1β production that is employed by DCs during their interaction with effector CD4 T cells. In the experiments above, we found that blocking Fas-FasL interaction led to a complete loss of detectable IL-1β in the supernatants, without affecting the synthesis of pro-IL-1β (Fig. 2b and 2c). Additionally, effector CD4 T cells were found to constitutively express FasL that was further upregulated upon their interaction with DCs (Supplementary Fig. 2c). These observations suggested the possibility that Fas signaling in DCs could be involved in the cleavage of pro-IL-1β. Indeed Fas deficient DCs from *lpr* mice^28^ were unable to secrete cleaved IL-1β underlining the necessity of DC intrinsic Fas signaling in T cell induced IL-1β production (Fig. 3c and 3d). It has been previously shown that Fas signaling in macrophages can trigger cleavage of pro-IL-1β in a caspase-8 dependent manner^29^, however the physiological importance of such IL-1β cleavage is not known. We found that DC-T cell interaction led to maturational cleavage of caspase-8 (Fig. 3e). While TNFR signaling has been reported to cause caspase-8 activation^30^, we discovered that only Fas, but not TNFR, signaling is critical for caspase-8 cleavage during cognate DC-T cell interaction (Fig. 3e). Inhibition of caspase-8 enzymatic activity by IETD resulted in abrogation of IL-1β cleavage implicating caspase-8 as the protease involved in mature IL-1β production by DCs (Fig. 3f). Caspase-8 has been shown to be important for induction NF-κB dependent genes upon TLR activation, including pro-IL-1β^31,32^. However, during DC-T cell interactions, caspase-8 inhibition did not affect pro-IL-1β synthesis (Fig. 3f). Furthermore, genetic deletion of caspase-8, along with RIPK3 to prevent necrosis^33^, in DCs led to a complete loss of secreted IL-1β (Fig. 3g). Although caspase-8 has been previously implicated in proteolytic cleavage of pro-IL-1β upon TLR4 acitvation^34^, our experiments demonstrate a crucial role for caspase-8 in IL-1β production by DCs in the absence of any PRR activation. Together, these data show that effector CD4 T cells can instruct IL-1β production by DCs thereby revealing a novel PRR-independent mechanism of innate immune activation.

**Figure 3.**
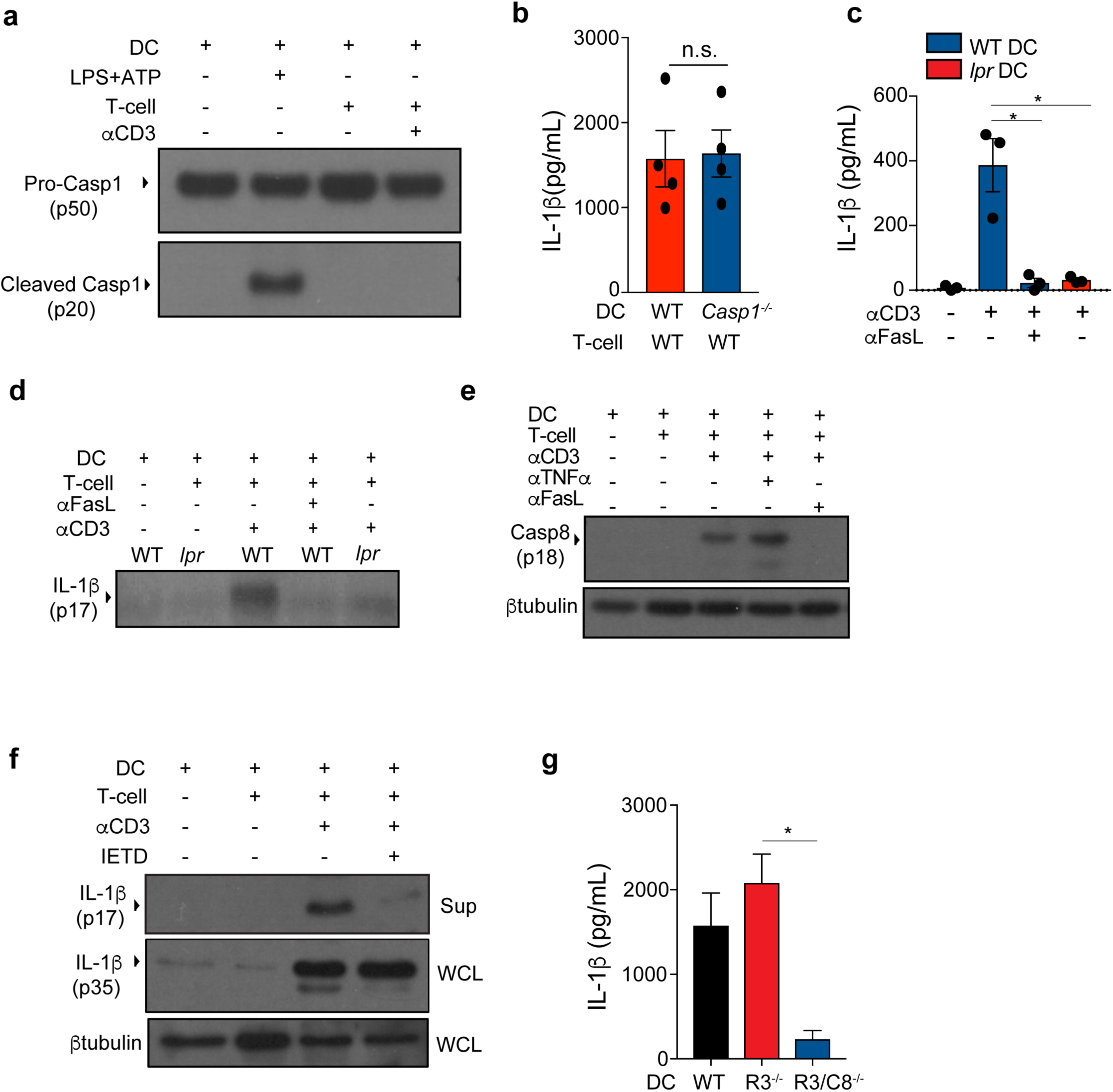
FasL-Fas interaction between effector CD4 T cells and DCs leads to caspase-8 dependent cleavage of pro-IL-1β. (a) WT DCs were stimulated with LPS+ATP or primed CD4 T cells (Th0). Cleaved caspase-1 was analyzed in the whole cell lysates 18h post stimulation (b) WT Th0 cells were stimulated with αCD3 in the presence of either WT or *Casp1^-/-^* DCs and supernatants were collected after 18h to measure IL-1β. (c, d) WT or *lpr* DCs were used to stimulate effector CD4 T cells (Th0) using αCD3 in the presence or absence of anti-FasL (10μgml^-1^) for 18h. Secreted cleaved IL-1β was quantified by (c) ELISA and (d) Western blot. (e) WT DCs were used to stimulate (αCD3) effector CD4 T cells (Th0) in the presence of anti-FasL (10μg ml^-1^) or anti-TNFα (20ug ml^-1^) for 12h. Caspase-8 activation was measured by assessing cleavage of caspase-8 in the WCL. (f) WT DCs were used to stimulate (using αCD3) effector CD4 T cells in the presence of IETD (10μM) for 18 h. Cleaved IL-1β (p17) and pro-IL-1β (p35) was measured by Western blot of culture supernatants and WCL, respectively. (g) WT, *Rip3^-/-^* (R3) and *Rip3*^-/-^*Casp8^-/-^* (R3/C8) DCs were used to stimulate effector CD4 T cells (Th0) in the presence of αCD3 for 18h and secreted IL-1β was quantified using ELISA. Error bars indicate SEM; (b, c and g) paired *t*-test, (a, d-f) Data are representative of two independent experiments.

Pathogen sensing by PRRs leads to inflammation primarily caused by secretion of pro-inflammatory cytokines and chemokines from myeloid cells^35^. Our data now suggest that effector CD4 T cells, irrespective of their specificity, could also induce inflammation by eliciting IL-1β production by DCs. Self-reactive CD4 T cells are key players in several IL-1β mediated autoimmune diseases such as RA, type 1 diabetes and multiple sclerosis (MS)^36–38^. However, whether self reactive CD4 T cells contribute to innate immune pathology by triggering IL-1β production has never been investigated. To test the ability of *bona fide* self-reactive CD4 T cells to elicit IL-1β production by interacting DCs, we utilized BDC2.5 TCR Tg diabetogenic mice^39^. BDC2.5 CD4 T cells are specific to a known islet cell peptide^40^. We stimulated these CD4 T cells *in vitro* using autologous DCs in the presence of their cognate peptide. Similar to the *in vitro* generated effector CD4 T cells, islet cell-reactive T cells also induced IL-1β production by DCs in a FasL and TNFα dependent manner (Fig. 4a).

**Figure 4.**
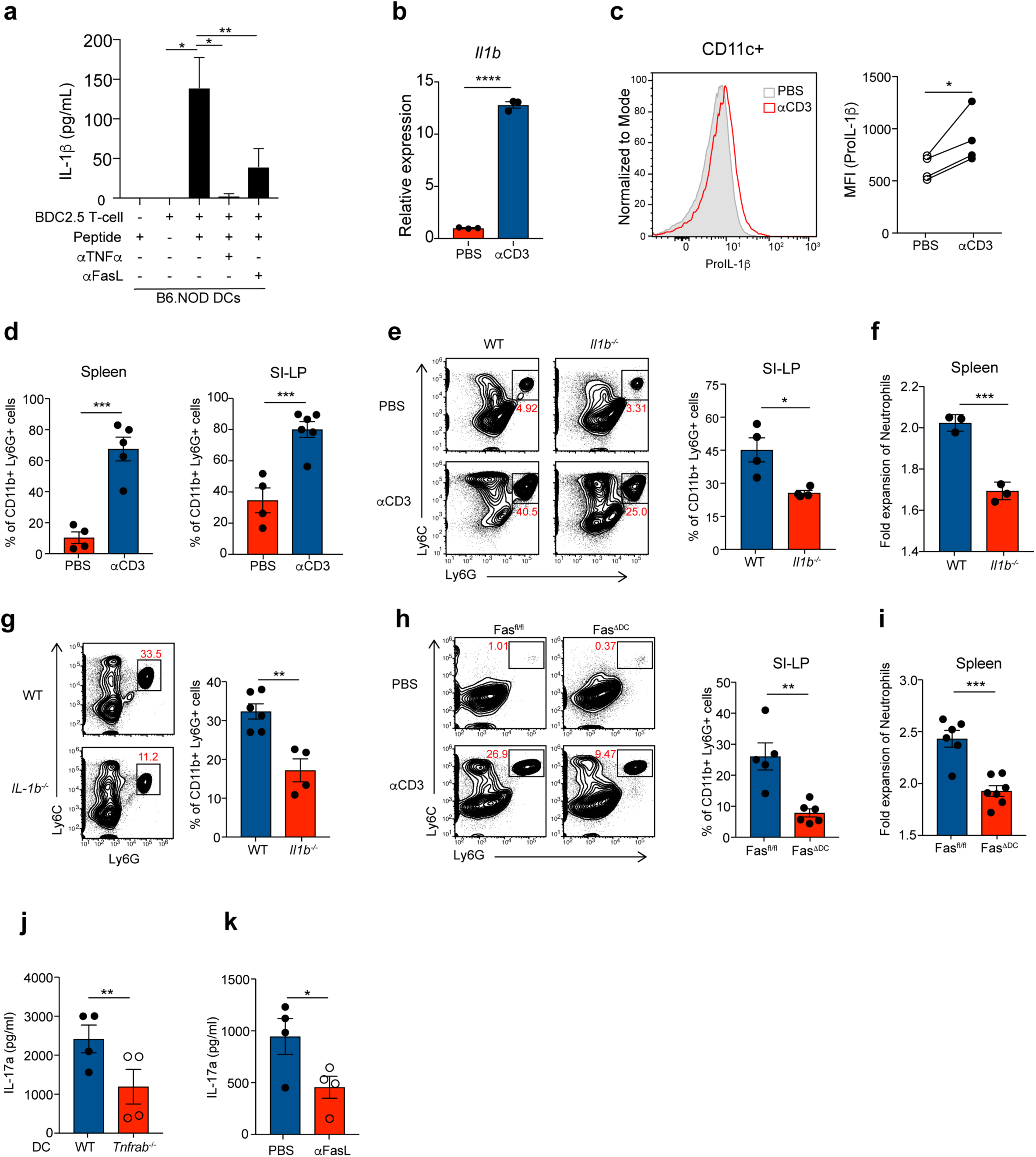
IL-1β produced upon DC-T cell cognate interaction leads to systemic inflammation marked by inflammatory leukocyte recruitment. (a) Total CD4 T cells were isolated from BDC2.5 TCR Tg mice and stimulated with B6.NOD BMDCs loaded with cognate peptide (50µM) for 18h and IL-1β was measured in the supernatant (b) WT mice were injected with αCD3 (50ug) intravenously. Spleen cells were harvested 3-4h post injection and immediately lysed for RNA isolation. Data are normalized to *Hprt1*. (c) Spleen cells from (b) were stained for DC markers and intracellular pro-IL-1β. Flow plot is pre-gated on live CD11c+ve cells. Data are representative of 2 independent experiments. (d) WT mice were injected with αCD3 (20μg) i.p. Neutrophil infiltration was quantified in the spleen (left panel) and SI-LP (right panel). (e and f) WT or *Il1b^-/-^* mice were treated with αCD3 i.p. (e) SI-LP and (f) spleens were harvested 18h later and analyzed for neutrophil infiltration. (g) WT OT-II T cells were differentiated into Th17 cells *in vitro*. 5 × 10^6^ OT-II Th17 cells were transferred i.v. into WT or *Il1b^-/-^* mice. 24h later 50µg of OVA_323-339_ peptide was injected i.v. into recipients. 12h after peptide injection, spleens were harvested and analyzed for neutrophil infiltration. (h,i) Fasfl/fl or Fasfl/fl x CD11c-cre (Fas^ΔDC^) mice were treated with αCD3 i.p.; 18h later (h) SI-LP and (i) spleen were analyzed for neutrophil infiltration. (j,k) CD44^hi^ CD62L^lo^ effector CD4 T cells were isolated from WT mice and re-stimulated with (j) WT or *Tnfrab^-/-^* CD11c+ve splenic DCs (k) WT DCs were used to stimulate CD44^hi^ CD62L^lo^ effector CD4 T cells in the presence or absence of anti-FasL neutralizing antibody and αCD3 (30ng/ml). (j,k) Quantities of IL-17a were measured in the culture supernatants 48h post stimulation. Error bars indicate SEM; (a-c, j,k) paired *t*-test, (d-i) unpaired *t*-test.

Although cytokines made by self-reactive T cells contribute to autoimmune inflammation, innate immune activation is known to be responsible for the precipitation of autoimmunity^41^. The pathology of these diseases is associated with infiltration of neutrophils and inflammatory monocytes into the affected tissues. Since the majority of existing mouse models of T cell driven autoimmunity still rely on initial PRR activation, usually in the form of *Mycobacterium tuberculosis*, to break tolerance^42^, we employed two new PRR independent approaches to mimic cognate DC-T cell interaction likely to occur during auto-immune flares. In the first approach, we employed *in vivo* administration of αCD3 speculating that systemic TCR activation would result in widespread T cell reactivation by myeloid cells. Intravenous administration of αCD3 led to rapid transcriptional upregulation of *Il1b* in the splenocytes (Fig. 4b). Upregulation of pro-IL-1β protein expression in CD11c+ DCs was also observed upon αCD3 stimulation (Fig. 4c). Moreover, we saw significant recruitment of neutrophils to the spleen as well as to the small intestinal lamina propria (SI-LP) within 18 hours of stimulation (Fig. 4d). More importantly, the inflammatory leukocyte recruitment in both the spleen and the SI-LP was significantly dependent on IL-1β (Fig. 4e, 4f and Supplementary Fig. 3a, b), establishing that *in vivo* interactions between T cells and myeloid cells leads to acute IL-1β production, which then promotes inflammatory leukocyte infiltration. In the second approach, we adoptively transferred OT-II TCR Tg Th17 T cells and induced their activation by intravenous administration of the MHCII-restricted OVA peptide (OVA_323-339_). Based on our *in vitro* data, we hypothesized that interaction of activated CD4 T cells with DCs bearing cognate peptides *in vivo* would lead to IL-1β production and result in IL-1β dependent inflammation as measured by expansion of circulating neutrophil populations. In agreement with our hypothesis, we observed that recruitment of neutrophils to the spleens of OT-II T cell receipient mice was dependent on IL-1β (Fig. 4g). Since we found that Fas signaling in DCs to be the most critical mediator of IL-1β production *in vitro*, we examined the requirement of Fas for T cell induced inflammation *in vivo*. We specifically ablated Fas in CD11c expressing cells (Supplementary Fig. 3c) and investigated its impact on IL-1β mediated leukocyte recruitment following DC-T cell interaction. Fas deletion in DCs significantly inhibited neutrophil infiltration in the SI-LP and spleen (Fig 4h and 4i) thus providing *in vivo* corroboration of our *in vitro* findings.

We have previously shown that IL-1β is a critical “licensing signal” for IL-17 production^6^. As we have found an important role for TNFα and Fas in IL-1β production by DCs, we reasoned that TNFR and Fas signaling in DCs may be necessary for cytokine production by Th17 cells during their reactivation. In agreement with our hypothesis, DCs lacking TNFR also had diminished capacity to trigger IL-17 production by effector CD4 T cells (Fig. 4j). Similarly, we discovered that blocking Fas signaling during T cell reactivation significantly compromised IL-17a production (Fig. 4k). We propose that this novel pathway of IL-1β production, mediated by TNFR and Fas signaling, evolved to enable optimal CD4 T cell effector function. However, cognate interactions of self-reactive effector T cells with antigen presenting myeloid cells are also likely to result in production of bioactive IL-1β, leading to detrimental immunopathology.

Our study thus establishes a previously unknown feature of DC-T cell cross-talk where effector CD4 T cells can instruct the innate immune system to produce IL-1β, independent of inflammasome activation. Of note, the signaling requirements of the “T cell instruction”, as described here, are parallel to the inflammasome pathway, in that it also relies on two independent signals that govern pro-IL-1β synthesis and its activation by proteolytic cleavage. The distinction between the two mechanisms of IL-1β production can be appreciated with regard to their physiological ramifications. The “TLR-NLR” inflammasome pathway is primarily employed by macrophages, resulting in production of IL-1β that is necessary for clearance of virulent pathogens (Supplementary Fig. 4). In contrast, the T cell instructed IL-1β, made by DCs as a result of TNFR-Fas signaling, is likely to be responsible for auto-immune flares in the absence of any overt pathogenic insult (Supplementary Fig. 4). Also, since the quantities of IL-1β produced by DCs are directly dictated by the concentration of the peptide, it is unlikely that bystander or low avidity interaction between DCs and T cells will trigger IL-1β production by DCs. Self-reactive CD4 T cells that escaped thymic selection and underwent effector differentiation perhaps employ this inflammasome-independent pathway of IL-1β production during cognate interaction with DCs, thereby causing innate inflammation associated with autoimmunity. Whether self reactive T cells in fact induce IL-1β using the “TNFR-Fas” pathway to elicit autoimmunity, requires further testing in T cell dependent autoimmune models.

Aberrant inflammasome activation caused by the gain-of-function mutations in NLRP3 and Pyrin drive IL-1β dependent auto-inflammatory conditions that are also initiated independently of TLRs and other PRRs^14^. The role of inflammasome in T cell driven autoimmune diseases, however, is neither discernible nor established. Strikingly, several lines of evidence support the involvement of TNFR and Fas signaling in autoimmune inflammation. Blocking of TNFα has been shown to effectively inhibit production of IL-1β by human synovial cells^43^. T cell intrinsic deletion of TNFα as well as Fas deficiency attenuates EAE disease severity^44,45^. Disruption of *Fas* also prevents autoimmune β cell destruction in non-obese diabetic mouse model^46^. Moreover, single nucleotide polymorphisms have been reported in the Tnfrsf1a, Faslg and Fas loci in strong association with IL-1β mediated autoimmune diseases^47–52^, thus providing credence to the argument that IL-1β in chronic autoimmune conditions could be dependent on “T cell instruction”, rather than inflammasome activation. Our data establishes a novel mechanism of IL-1β production that is initiated by effector CD4 T cells and thus presents an explanation for the presence of IL-1β seen in T cell mediated autoimmunty.

## Acknowledgements

We thank all the members of the Pasare lab for helpful discussions. Thanks to Rustam Bagirzadeh and Margaret McDaniel for critical reading of the manuscript. This work was supported by grants from the National Institutes of Health (AI113125 and AI123176) to C.P.

## Author contributions

C.P. and A.J. conceptualized the study, designed the experiments and wrote the manuscript. A.J. performed majority of the experiments. R.A.I.C. and A.S.C. performed some of the experiments. J.K. and K.C. helped with BDC2.5 TCR Tg mouse experiments. A.C. provided Fas floxed mice. A.O. helped with *Rip3*^-/-^ and *Rip3^-/-^Casp8^-/-^* mice experiments. N.P. helped with *Rip3^-/-^Casp8^-/-^* dendritic cell experiments.

## Methods

### Mice

C57BL/6 wild-type control mice were obtained from the UT Southwestern Mouse Breeding Core Facility. IL-1β ^-/-^ mice were generated by David Chaplin, UA at Birmingham and provided to us by Fayyaz S. Sutterwala at Cedars Sinai. Fas floxed mice were provided to us by Alexander Chervonsky at University of Chicago. BDC2.5 Tg C57Bl6/-I-A^g7^ mice were provided by Jonathan Katz at Cincinnati Children’s Hospital Medical Center. RIP3^-/-^ and RIP3^-/-^Caspase8^-/-^ KO were provided by Andrew Oberst at the University of Washington. B6.MRL-*Fas^lpr^*/J (*lpr*), B6.129S-*Tnf^tm1Gkl^*/J (Tnfa^-/-^), B6.129S-*Tnfrsf1a^tm1Imx^ Tnfrsf1b^tm1Imx^*/J (Tnfrab^-/-^) were obtained from Jackson Laboratories. All mice were bred and housed in a specific pathogen-free facility at UT Southwestern Medical Center. For isolation of steady state CD4 memory T cells, mice were housed in a conventional facility for 2-4 weeks before tissue isolation. All mouse experiments were done as per protocols approved by Institutional Animal Care and Use Committee (IACUC) at UT Southwestern Medical Center and Cincinnati Children’s Hospital Medical Center.

### Reagents

IL-1Ra (R&D Systems; 480-RM, 50ng/ml), rTNFα (Peprotech; 315-01Am, 20ng/ml), αCD3e (Biolegend; 100331 for *in vitro* assays and 100340 for *in vivo assays*), αTNF (Biolegend; 506332, 20µg/ml), αFasL (Biolegend; 106608, 10µg/ml), αCD40L (Biolegend; 310812, 10 µg/ml), IL-1β ELISA (R&D Systems; DY-401), TNFα ELISA(Biolegend; 510802 for capture and 506312 for detection), Naïve T cell Mojosort (Biolegend; 480040), total CD4 T cell Mojosort (Biolegend; 480006), Zombie yellow (Biolegend; 423103), DAPI (Biolegend; 422801), IETD (R&D Systems; FMK007, 10µM), rGMCSF (Biolegend; 576306), Anti-biotin beads (Milteny Biotec; 130-090-485), OVA_323-339_ (Invivogen; vac-isq), LPS (Sigma, 100ng/ml), ATP (Invivogen; tlrl-atpl, 5nM), IL17a ELISA (Biolegend; 506902 for capture and 507002 for detection), BDC2.5 peptide (RTRPLWVRME, custom ordered from Synthetic Biomolecules)

## Antibodies for flow cytometry

Anti-mouse Pro-IL-1β APC (eBioscience, 17-7114-80; 1:500), Anti-mouse CD11b BV785 (Biolegend, 101243; 1:400), Anti-mouse CD11C FITC (Biolegend; 117306, 1:400), Anti-mouse Ly6G FITC (Biolegend, 127605; 1:400), Ly6C BV711 (Biolegend, 12803; 1:400), Anti-mouse TNFα FITC (Biolegend, 506304, 1:1000), Anti-mouse F4/80 APC-eflour 780 (Invitrogen, 47-4801-80; 1:400), Anti-mouse CD90 Pacific blue (Biolegend, 105324; 1:400), Anti-mouse FasL PE (Biolegend, 106605; 1:100), Anti-mouse CD62L biotin (Biolegend, 104404; 1:200), Anti-mouse CD11c biotin (Biolegend,117304; 1:500)

### Cell culture

Complete RPMI media (RPMI1640 (Hyclone) supplemented with L-glutamine, penicillin-streptomycin, sodium pyruvate, β-mercaptoethanol (Sigma)) was used throughout the experiments.

### In vitro differentiation of naïve CD4 T cells

Cell culture treated plates were coated with αCD3 (5μg mL^-1^) and αCD28 (5μg mL^-1^) for 3-4hrs at 37ºC. CD4^+^CD62L^hi^ naïve CD4 T cells were isolated from splenocytes using a Mojosort naïve CD4 T cell isolation kit according to manufacturer’s protocol. Purified naïve T cells were plated in antibody-coated plates with appropriate polarizing conditions for 5 days. T cell were cultured in complete RPMI supplemented with 10% FCS. Cytokine cocktails for *in vitro* polarization: Th1 – IL-12 (10ng mL^-1^, peprotech), IL-2 (50U mL^-1^, peprotech), αIL-4 (10μg mL^-1^, Biolegend); Th17 – IL-6 (20ng mL^-1^, peprotech), hTGFβ (5ng mL^-1^, peprotech), αIL-4 (10μg mL^-1^, Biolegend), αIFNγ (10μg mL^-1^, Biolegend), IL-23 (20ng mL^-1^, Biolegend) and IL-1β (20ng mL^-1^, peprotech), Th2-IL-4 (4ng mL^-1^, peprotech), IL-2 (50U mL^-1^, Biolegend), and αIFNγ (10μg mL^-1^, Biolegend), Th0 – IL-2 (50U ml^-1^). For co-culture experiments T cells were rested in 10% RPMI supplemented with IL-2 (10U mL^-1^) for 36hrs before co-culture.

### In vitro differentiation of bone marrow derived dendritic cells and retroviral transduction

Mouse progenitors were isolated from bone marrow (femurs and tibias). Following RBC lysis, cells were plated at 0.75 × 10^6^ mL^-1^ in BMDC media (5% FCS containing complete RPMI + 1% rGMCSF (Biolegend, 100ng mL^-1^)). Media was replaced on day 2 and day 4 and cells were harvested for experiments on day 5 by gently flushing each well. For retroviral transduction, following RBC lysis cells were plated at 10 × 10^6^ mL^-1^ in 2mL of retroviral supernatant containing 8µL mL^-1^ polybrene. Cells were spinfected at 2500 rpm, 32ºC, for 90 minutes, then 3mL of BMDC media was added to each well and cells were incubated overnight. The next morning, ~70% of the media was removed from the wells and spinfection was repeated with fresh retroviral supernatant. On day 5 cells were harvested for experiments.

### Co-culture of DC and T cells

1 million DCs were cultured with 4 million T cells in 12 well plate. DC-T cell interaction was triggered by adding either αCD3e (αCD3, 200ng mL^-1^) or OVA_323-339_ as described in the legends. In experiments measuring secreted IL-1β, IL-1R antagonist (50ng mL^-1^) was added 1hr prior to DC stimulation to block IL-1β consumption. Co-cultures were also pretreated with neutralizing antibodies and inhibitor wherever described. Co-culture experiments were performed in complete RPMI supplemented with 10% FCS. For Western blot analysis of the culture supernatant cells were cultured in 1% FCS containing complete RPMI.

### T cell re-stimulation (*in vivo*)

6-8 weeks old mice were treated with 20μg αCD3 or PBS by i.p. injection. Lamina propria cells and splenocytes were isolated at given time points after stimulation followed by surface and intracellular staining.

### Quantitative real-time PCR

RNA was isolated using Qiagen RNA extraction kit using manufacturer’s protocol. cDNA was synthesized using Random primers and MMLV reverse transcriptase (Invitrogen; 2805-013). The QuantStudio 7 Flex Real-Time PCR System (ThermoFisher Scientific) was used to measure SyBR green (ThermoFisher Scientific) incorporation. All data is normalized to 18s. qPCR primers sequences are as follows: *Il1b*: Fwd-5’ TGTGCTCTGCTTGTGAGGTGCTG 3’, Rev 5’CCCTGCAGCTGGAGAGTGTGGA3’ | *Hprt1*:Fwd-5’ CAGTCCCAGCGTCGTGATTA-3’, Rev-5’ TGGCCTCCCATCTCCTTCAT-3’ | *18S*: Fwd-5’ GTAACCCGTTGAACCCCATT, Rev-5’ CCATCCAATCGGTAGTAGCG

### Western blot analysis

Cells were lysed in 1X RIPA Buffer and protein was quantified using Pierce™ BCA Protein Assay Kit. Cell lysates where boiled in 1X Laemmli buffer at 95ºC for 10 mins. Cell lysates were separated by SDS–PAGE and transferred onto PVDF membranes. Blots were incubated with anti-IL1β [1:1000] (R&D AF-401-SP), anti-caspase8 [1:1000] (Enzo ALX-804-447-C100), anti-caspase1 [1:1000] (Genentech), anti-β tubulin [1:5000] (CST 2146S). As secondary antibodies, anti-rabbit-IgG-HRP (Biorad) [1:5000], anti-mouse-IgG-HRP and anti-goat-IgG-HRP [1:10000] (Jackson ImmunoResearch Laboratory) were used. Anti-β-actin (C4, Santa Cruz, 1:5000) was used as control. Western blot was develop using SuperSignal™ West Pico PLUS Chemiluminescent Substrate (Thermo Fisher) and ECL signal was recorded on X-Ray Films using a developer (Kodak).

### Surface and intracellular staining and flow cytometry

Cells were stained with relevant antibodies for 30 min on ice and washed. For intracellular staining, Foxp3 staining buffer set (eBioscience; 00-5523-00) was used according to manufacturer’s protocol. The stained cells were analyzed with BDLSRII or Novocyte (ACEA biosciences). For cytokine receptor staining, control refers to fluorescence minus one control. Data were analyzed with FlowJo 10 Software.

### Isolation of Lamina Propria Lymphocytes

Small intestines were flushed with cold PBS and carefully cut longitudinally. 1-2cm pieces of the tissue were digested twice with 2mM EDTA buffer followed by 3 rounds of enzymatic digestion with Collagenase IV (10 µg/ml, Sigma) and DNase I (500 µg/ml, Sigma). Cell suspensions obtained after digestions were loaded on a 40-70% percoll gradient as described before.

### Quantification and statistical analysis

Based on previous and preliminary studies in our lab, we predicted that the reported samples sizes would be sufficient to ensure adequate power. Statistical analyses were performed in Prism (Graph pad) using unpaired or paired *Student’s t test* as indicated in the figure legends. Data are presented as means [Symbol] SEM. Significance was considered at *\*p*<0.05, **p<0.01, ***p<0.001, ****p<0.0001. n.s. = not significant.

### Data availability statement

The data generated during and/or analyzed during the current study are available from the corresponding author on reasonable request.

