## Supplementary material for "T cells instruct dendritic cells to produce inflammasome independent IL-1β causing systemic inflammation"

### Supplementary Figure 1

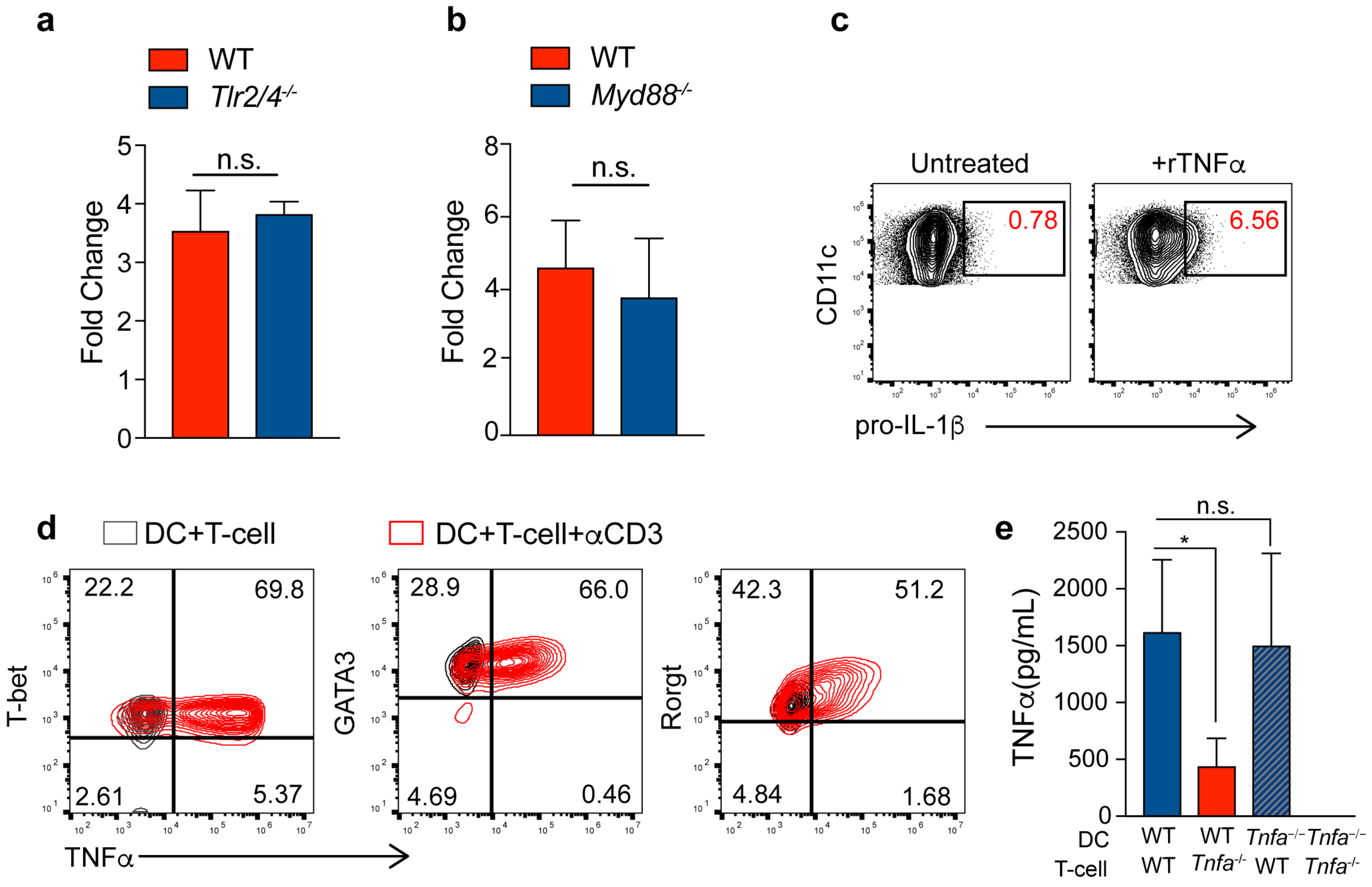

### Supplementary Figure 2

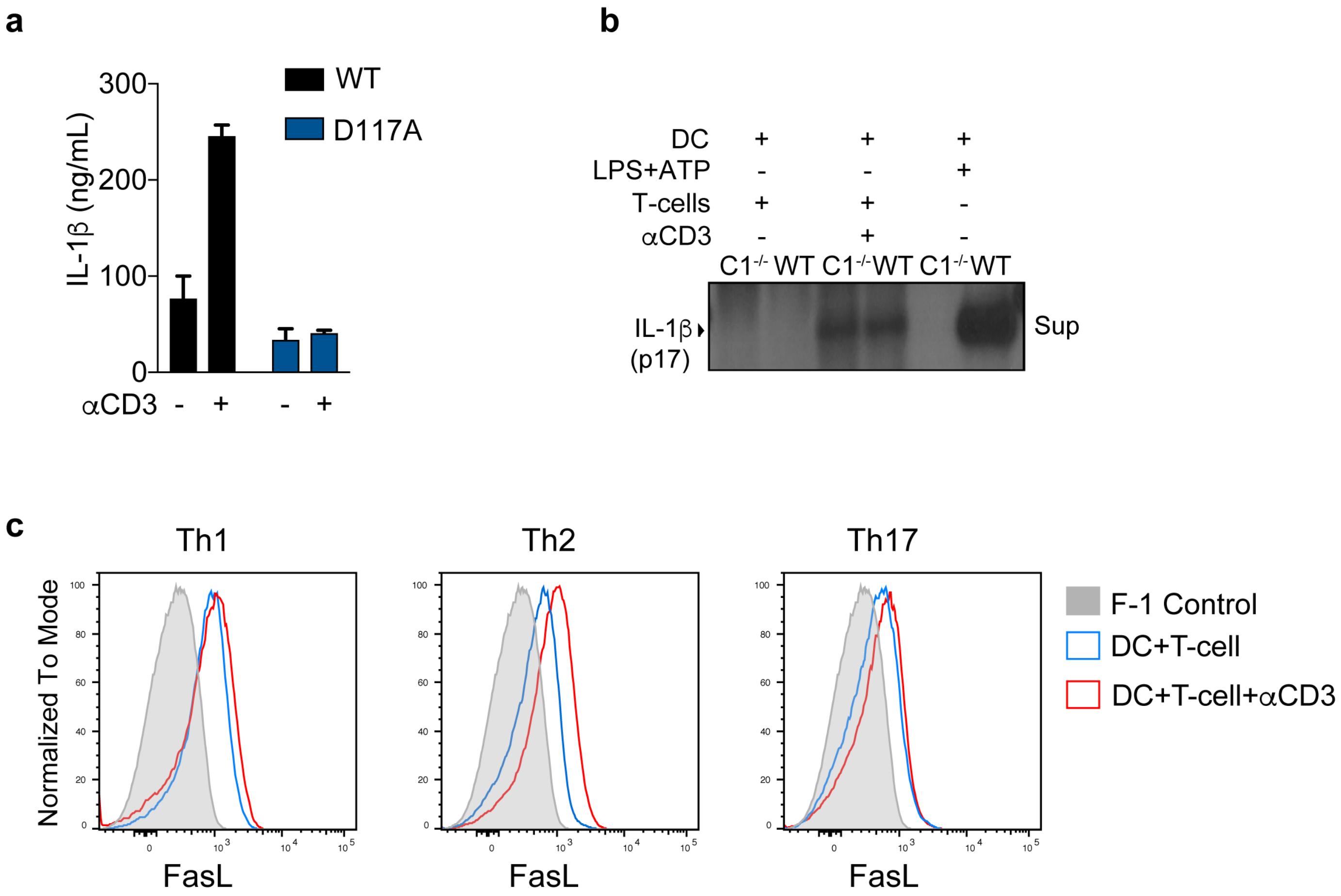

### Supplementary Figure 3

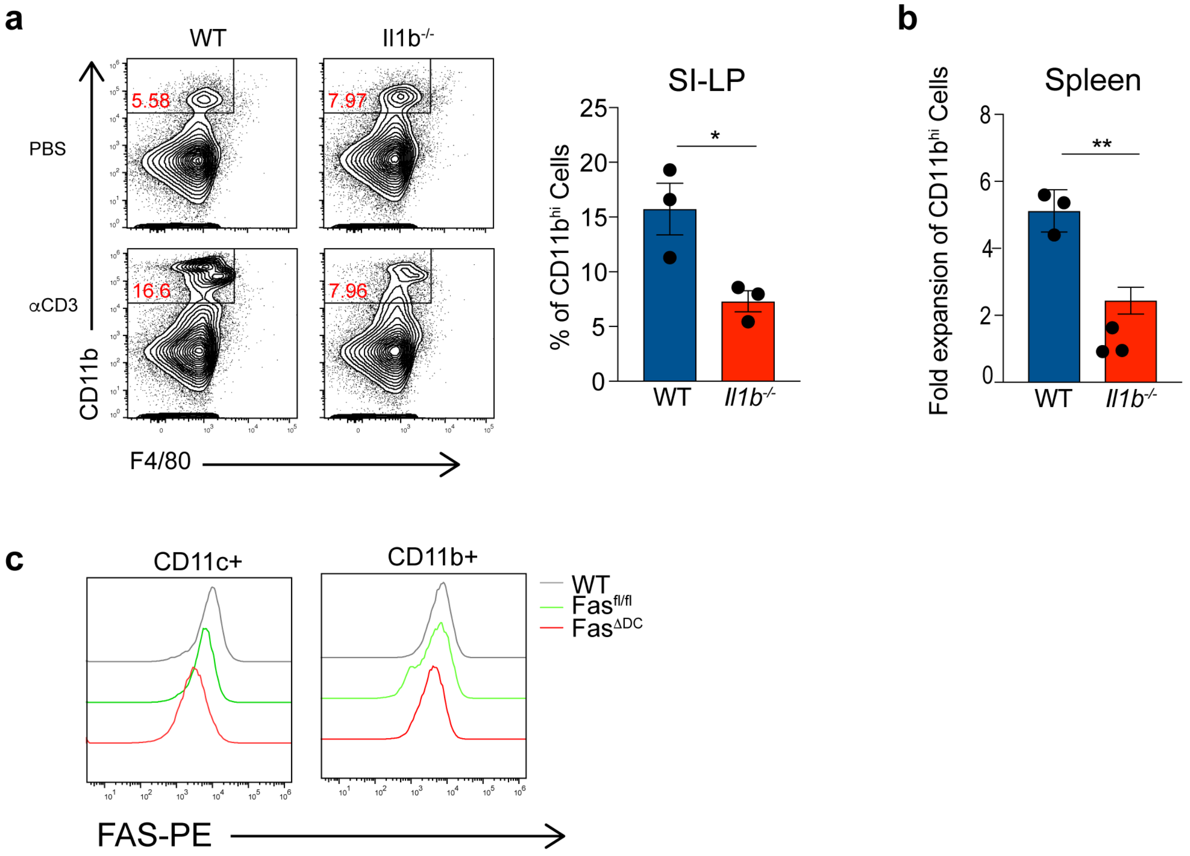

### Supplementary Figure 4

**a**

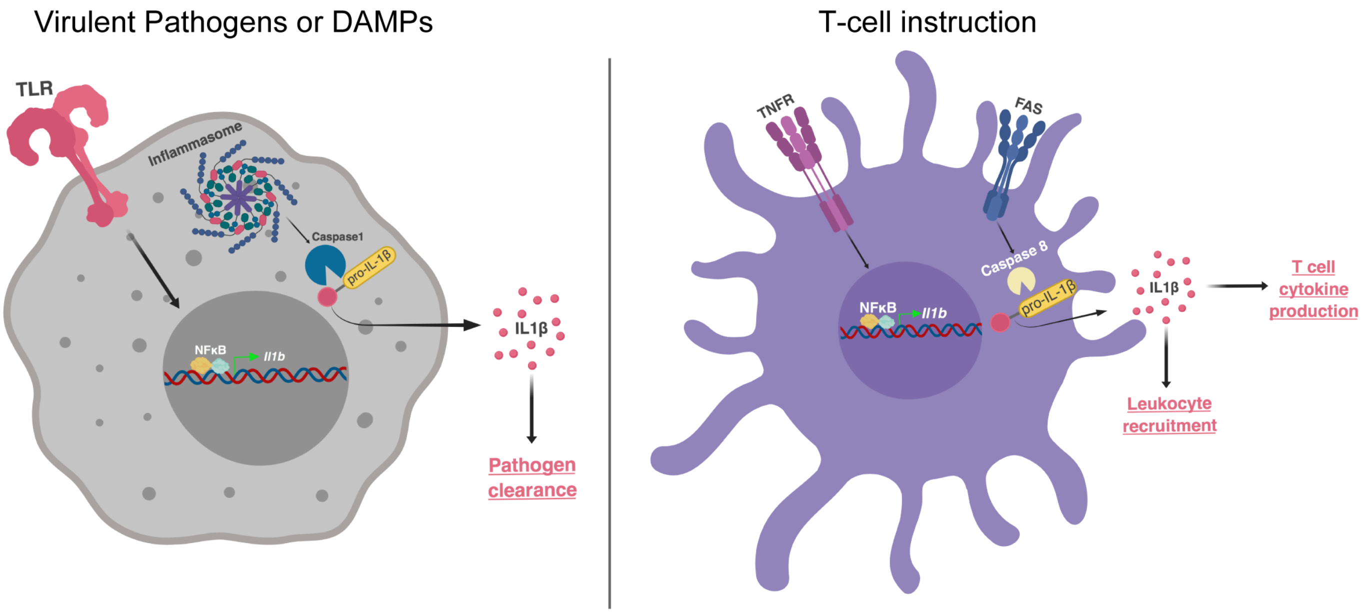

#### Supplementary Figure Legends

**Supplementary Figure 1. Induction of pro-IL-1β in DCs is independent of TLR activation but dependent on TNFα.**

(a and b) WT, *Tlr2/4*$^{-/-}$ or *Myd88*$^{-/-}$ DCs were stimulated with WT Th0 cells using αCD3. Intracellular pro-IL-1β was analyzed 6h post stimulation. Fold change indicates proportion of pro-IL-1β+ve DCs, compared to PBS controls (c) DCs were stimulated *in vitro* using recombinant TNFα (20ng/ml) for 6h followed by intracellular pro-IL-1β stain. Data are representative of two independent experiments. (d) Th1, Th2 and Th17 polarized CD4 T cells were stimulated with αCD3 using WT DCs in the presence of brefeldin A for 3h. Intracellular TNFα in T cells was analyzed using flow cytometry. Cells were considered to be transcription factor positive based on Isotype control antibody staining. Data are representative of three independent experiments. (e) WT or *Tnfa*$^{-/-}$ T cells were stimulated with WT or *Tnfa*$^{-/-}$ DCs. Supernatants from cultures of indicated genotypes (all in the presence of αCD3) were assayed for presence of TNFα 6h post stimulation. Error bars indicate SEM. (a, b and e) paired *t*-test.

**Supplementary Figure 2. Effector CD4 T cell induce Casapse-1 independent IL-1β production upon interaction with DCs.**

(a) *Il1b*$^{-/-}$ bone marrow precursors were transduced with MSCV construct expressing WT or D117A *Il1b* cDNA and differentiated into DCs *in vitro*. Transduced DCs were used to activate *in vitro* primed CD4 T cells (Th0) in the presence of αCD3. Secreted IL-1β was measured 18h post stimulation. Error bars indicate SEM. (b) WT or *Casp1*$^{-/-}$ DCs were used to stimulate CD4 T cells (Th0) using αCD3. IL-1β was analyzed in the culture supernatants 18 hours post stimulation. (c) *In vitro* primed Th1, Th2 and Th17 cells were co-cultured with WT DCs in the presence of αCD3. Surface FasL staining on live CD90.2+ve cells is shown with or without αCD3 stimulation. Data are representative of two independent experiments.

**Supplementary Figure 3. DC-T cell interaction *in vivo* leads to IL-1β dependent inflammatory cell recruitment.**

(a) WT and *Il1b*$^{-/-}$ mice were injected with αCD3 (20μg) i.p. and monocyte infiltration was analyzed in SI-LP, 18h later, using flow cytometry. (b) Splenocytes from mice in (a) were also analyzed for CD11b+ve cell recruitment. Data are plotted as fold change when compared to PBS control. (c) Fas expression on DCs from given genotypes shows CD11c+ cell specific Fas deletion. Error bars indicate SEM. (a and b) unpaired *t*-test.

**Supplementary Figure 4. Illustration of "T cell instructed" IL-1β production by DCs and its comparison to inflammasome induced IL-1β production by macrophages.**

(Left) During inflammasome activation in macrophages, TLR and NLR-Caspase-1 activation leads to synthesis and cleavage of pro-IL-1β, respectively. Robust production of IL-1β as a result of inflammasome activation is critical for pathogen clearance. (Right) In contrast, "T cell instructed" IL-1β production by DCs utilizes TNFR signaling for pro-IL1β synthesis while the cleavage signal is provided by Fas-Caspase-8 axis. The IL-1β produced upon T cell instruction drives cytokine production by effector CD4 T cells and systemic leukocyte recruitment, a hallmark of auto-immune pathology. Figure was created with BioRender.
